# Ca^2+^ ions promote fusion of Middle East Respiratory Syndrome coronavirus with host cells and increase infectivity

**DOI:** 10.1101/2019.12.18.881391

**Authors:** Marco R. Straus, Tiffany Tang, Alex L. Lai, Annkatrin Flegel, Miya Bidon, Jack H. Freed, Susan Daniel, Gary R. Whittaker

**Author notes:** Corresponding authors: Gary Whittaker, Susan Daniel. Both authors contributed equally to this work (Marco R. Straus, Tiffany Tang).

## Abstract

Middle East respiratory syndrome coronavirus (MERS-CoV) is a major emerging zoonotic infectious disease. Since its first outbreak in 2012, the virus has repeatedly transmitted from camels to humans with 2,468 confirmed cases, causing 851 deaths. To date, there are no efficacious drugs and vaccines against MERS-CoV, increasing its potential to cause a public health emergency. A critical step in the life cycle of MERS-CoV is the fusion with the host cell with its spike (S) protein as main determinant of viral entry. Proteolytic cleavage of S exposes its fusion peptide (FP), which initiates membrane fusion. Previous studies on the related severe acute respiratory syndrome coronavirus (SARS-CoV) FP have shown that calcium (Ca^2+^) plays an important role for fusogenic activity via a Ca^2+^ binding pocket with conserved glutamic acid (E) and aspartic acid (D) residues. SARS-CoV and MERS-CoV FP share a high sequence homology and here, we investigated whether Ca^2+^ is required for MERS-CoV fusion by substituting E and D residues in the MERS-CoV FP with neutrally charged alanines. Upon verifying mutant cell surface expression and proteolytic cleavage, we tested the mutants ability to mediate infection of pseudo-particles (PPs) on host cells without and with Ca^2+^. Our results demonstrate that intracellular Ca^2+^ enhances MERS-CoV WT PPs infection by approximately two-fold and that E891 is a crucial residue for Ca^2+^ interaction. Electron spin resonance revealed that this enhancement could be attributed to Ca^2+^ increasing MERS-CoV FP fusion-relevant membrane ordering. Intriguingly, isothermal calorimetry titration showed that MERS-CoV FP binds one Ca^2+^, as opposed to SARS-CoV FP which binds two. Our data suggests that there are significant differences in FP-Ca^2+^ interactions of MERS-CoV and SARS-CoV FP despite their high sequence similarity and that the number of Ca^2+^ ions interacting with the FP has implications on the fusion dynamics of the virus.

## Introduction

Coronaviruses (CoVs) comprise a family of enveloped viruses causing respiratory and/or enteric tract infections across a variety of hosts. Recently, the emergence of Middle East respiratory syndrome coronavirus (MERS-CoV), first isolated in 2012 from a 60-year-old man in the Kingdom of Saudi Arabia, has highlighted the pathogenic potential of CoVs [1]. To date, MERS-CoV has caused 2,468 laboratory-confirmed human infections worldwide, resulting in 851 deaths (case-fatality: 34.48%) [2]. However, human MERS-CoV originated in bats from where it transitioned into intermediate hosts such as dromedary camels[3–5]. It is currently believed that disease outbreaks in humans result from interactions with dromedary camels. Although transmissions between humans is rare, this route was responsible for a widespread outbreak of MERS-CoV in 2015 in South Korea [6]. Moreover, it was demonstrated that MERS-like bat CoV could potentially infect human tissues in the absence of an intermediate host when the respective tissue expresses a protease that is able to prime the virus for fusion [7]. More recently, the World Health Organization (WHO) listed MERS-CoV as one of ten diseases that requires urgent accelerated research and development because of its potential to cause a public health emergency with few or no efficacious drugs and/or vaccines to counter them [8]. To address these needs, it is important to have a more nuanced understanding of MERS-CoV infection process to develop therapeutics to target crucial steps in MERS-CoV infection.

MERS-CoV is an enveloped virus that utilizes a class I fusion protein for viral entry into the host cell [9]. The MERS-CoV spike (S) protein contains a receptor binding subunit (S1) and a membrane fusion subunit (S2), which facilitates binding to the host cell surface receptor, dipeptidyl peptidase 4 (DPP4), and fusion with the host cell membrane, respectively [10,11]. To mediate fusion with the host cell membrane, S needs to be cleaved by either host cell surface proteases or endosomal proteases (such as transmembrane protease serine 2 (TMPRSS2), furin, or cathepsins) [12–16]. Cleavage can occur at two different sites: the S1 and S2 boundary (S1/S2), and adjacent to a fusion peptide (FP) within the S2 domain (S2’) [17,18]. It is suggested that both cleavage events act in concert to promote membrane fusion and viral infectivity. The first cleavage event takes place at the S1/S2 site, which then is believed to open up the S2’ site for further proteolytic processing, resulting in the exposure of the FP. As a consequence, the FP inserts into the host membrane, mediating membrane fusion [19].

MERS-CoV viral fusion can occur at two different locations in the host cell: at the plasma membrane or at the endosomal membrane. Host cell surface proteases, such as TMPRSS2, can cleave MERS-CoV S for entry directly at the plasma membrane [13]. In the absence of cell surface proteases, MERS-CoV is endocytosed, and endosomal proteases, such as cathepsins, can cleave S for endosomal entry. [19]. However, in contrast to other viruses such as influenza A, MERS-CoV fusion does not seem to depend on a pH change in the endosomes, as receptor binding in conjunction with proteolytic activation are sufficient to trigger fusion [20]. A more thorough description of the fusion pathways are summarized in several excellent reviews [20–23].

A recent study on the FP of severe acute respiratory syndrome coronavirus (SARS-CoV) suggested that the FP region begins immediately downstream of the S2’ cleavage site and that it can be separated into two distinct domains, FP1 and FP2 [24]. The authors showed that both FP1 and FP2 increase membrane ordering and proposed that FP1 and FP2 together form an extended FP which acts as a bipartite fusion platform. Intriguingly, both FP subdomains required calcium cations (Ca^2+^) to exert their function in membrane entry and fusion. It was proposed that negatively charged aspartic acid (D) and glutamic acid (E) residues coordinate with the positively charged Ca^2+^ within the fusion platform to promote greater membrane ordering. Furthermore, the authors showed that depletion of extracellular as well as intracellular Ca^2+^ pools resulted in significantly reduced infectivity of SARS-CoV pseudo-particles, suggesting that both the plasma membrane and endosomal cell entry pathways are regulated by Ca^2+^. Other studies too have revealed an important role for Ca^2+^ in the fusion dynamics of enveloped viruses. Rubella virus (RuV), for example, requires Ca^2+^ for virus entry and infection [25]. Ca^2+^ is coordinated in between the two fusion loops of the RuV E1 fusion protein by asparagine (N) and aspartic acid and both residues were shown to be essential for virus viability. More recently, Ca^2+^ was reported to enhance the entry of Ebola virus (EBOV) by directly interacting with the FP of the Ebola virus fusion protein [26]. The authors were able to identify two negatively charged residues (D522 and E540) in the FP that interact with Ca^2+^ and this interaction is critical for viral fusion with the host cell.

Here, we addressed the question whether Ca^2+^ is required for viral entry and infectivity of MERS-CoV by creating a set of S protein mutants in which negatively charged residues in the FP were mutated to alanines (A). We used the different mutants in pseudo-particle infection assays to demonstrate that fusion of MERS-CoV is promoted but not dependent on Ca^2+^. Biophysical analysis demonstrated that Ca^2+^ interactions with the MERS-CoV increases membrane ordering in a Ca^2+^-dependent manner. Our data suggests that a single glutamic acid residue in a region complementary to the SARS-CoV FP1 domain is involved in Ca^2+^ coordination.

## Results

### S protein mutants with amino acid substitutions in the FP are expressed and cleaved as WT protein

In order to test if Ca^2+^ is required and which residues within the MERS-CoV FP are necessary for its coordination, we generated a set of mutated S proteins in which we substituted negatively charged aspartic acid and glutamic acid residues in the FP with alanine (A) (Figure 1A). We defined the FP region based on a sequence comparison with the SARS-CoV FP that was described previously [24] (Figure 1A). The S mutant plasmids were generated via site-directed mutagenesis and verified by sequencing (data not shown). We next tested whether the mutated S proteins were expressed by transfecting the plasmid DNA into HEK 293T cells. Since the MERS-CoV S protein can be cleaved by constitutively expressed proteases during the maturation process, we added a protease inhibitor, dec-RVKR-CMK, immediately after transfection to block this cleavage. (Figure 2A). Dec-RVKR-CMK was shown to inhibit a wide variety of proteases such as furin, cathepsins, trypsin, and TMPRSS2 to block S cleavage [15,27]. As we wanted to confirm that the mutant S proteins are properly expressed and trafficked to the cell membrane surface, like the WT S protein, we isolated proteins 18 h post transfection via cell-surface biotinylation. Western blot analysis of the cell surface proteins showed that all mutant S proteins were expressed to similar levels as the MERS-CoV WT S protein (Figure 2B). With all mutant S proteins expressed, we next wanted to investigate whether the mutants exhibit the same cleavage pattern as WT S since proteolytic cleavage is an important and crucial trigger for viral fusion. We accomplished this by treating transfected cells with trypsin prior to protein isolation. Western blot analysis revealed that the cleavage pattern of the mutated S proteins was similar to that of WT S, suggesting that the introduced mutations in the FP did not impair protease accessibility and cleavage (Figure 2B).

**Figure 1:**
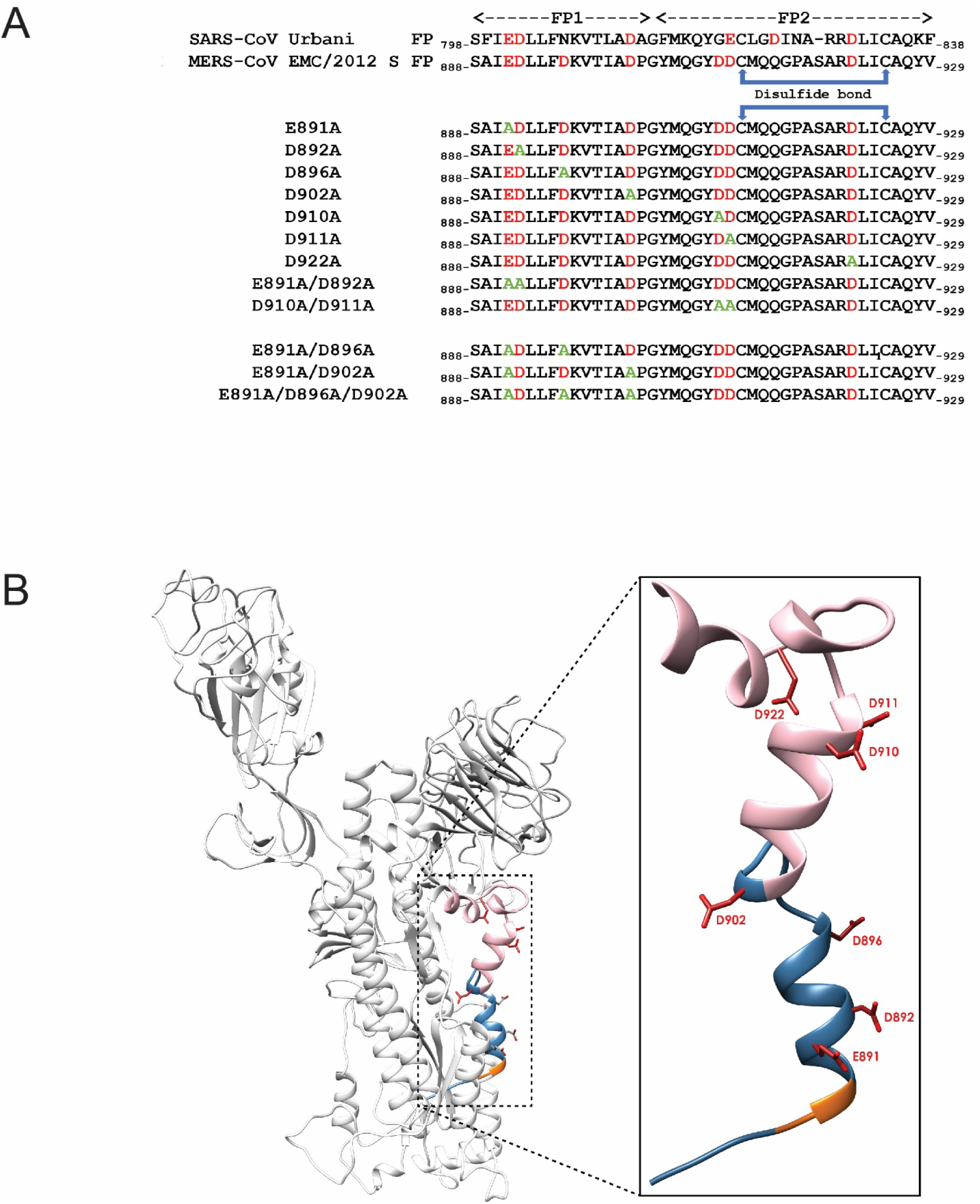
Sequence and model of MERS-CoV S fusion loop. (A) Sequences of SARS-CoV S Urbani and MERS-CoV S EMC/2012 fusion peptides (FP). FP1 and FP2 designate the two different domains in the FP as described previously (Lai Millet). Sequences below illustrate the mutations that were introduced in the MERS-CoV S protein via site-directed mutagenesis. In red are the negatively charged residues D and E, in green are the A substitutions. (B) Modeling of the MERS-CoV S monomer with an emphasis on the FP. Negatively charges Ds and E are depicted as atomic bonds in red. The S2’ site is orange and the FP1 and FP2 domains are labeled blue and pink, respectively.

**Figure 2:**
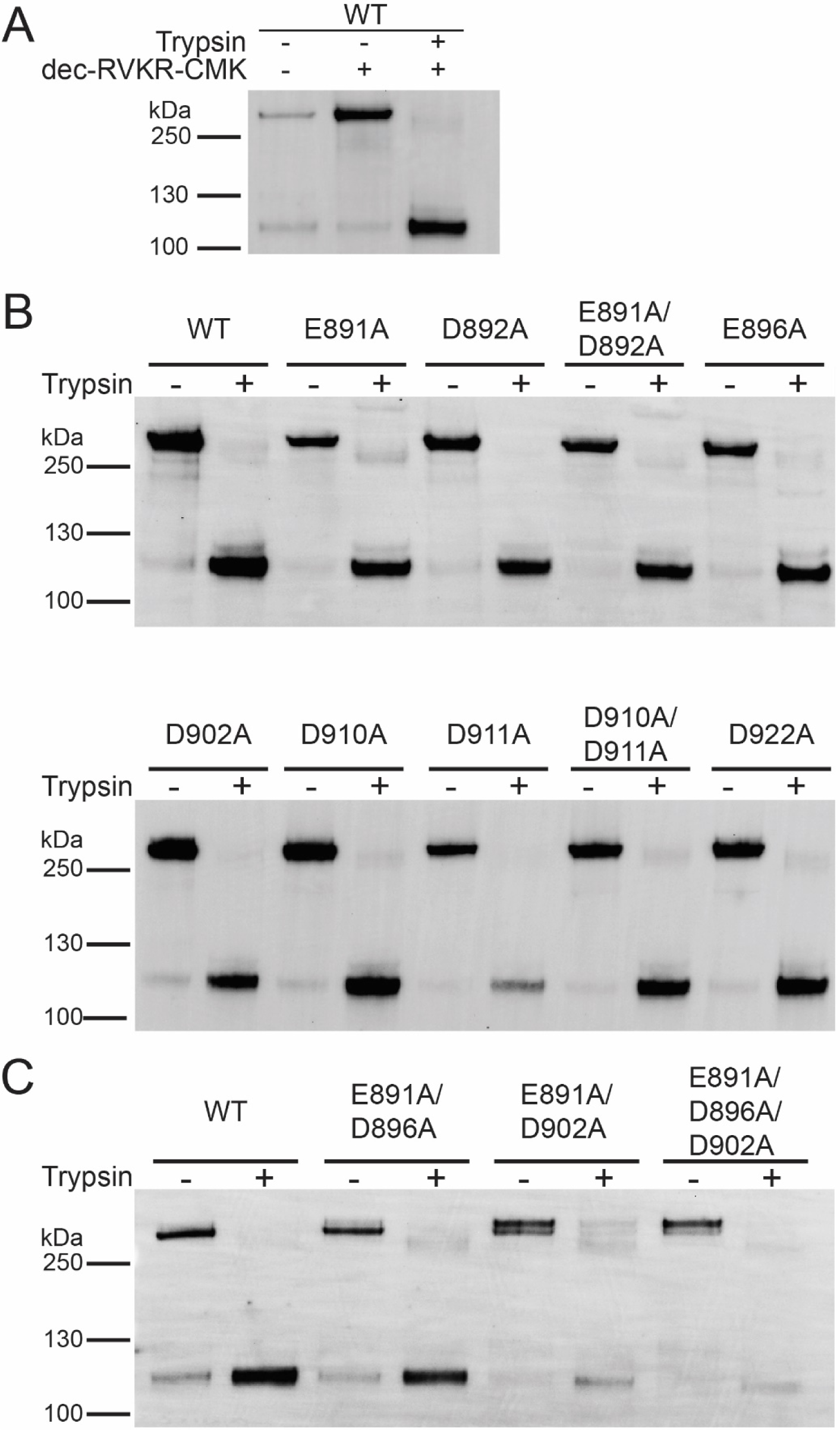
Protein expression and trypsin-mediated cleavage of MERS-CoV S WT and mutants. (A) Plasmid DNA encoding MERS-CoV S WT EMC/2012 was transfected in HEK293T cells. The protease inhibitor dec-RVKR-CMK at a concentration of 75 µM was added at the time of transfection, as indicated. After 18 h, transfected cells were treated with 0.8 nM TPCK-treated trypsin, as indicated. Proteins were subsequently isolated via cell-surface biotinylation. The cell surface proteins were analyzed using SDS-PAGE and detected using a Western blot with MERS-CoV S antibodies. (B) and (C) MERS-CoV S mutant proteins with indicated A substitutions were expressed in HEK293T cells. Protease inhibitor dec-RVKR-CMK was added at the time of transfection and after 18 h, cells were treated with TPCK-treated trypsin, as indicated. Cell surface proteins were isolated and analyzed as described above. Full length S proteins are visible at approx. 250 kDa. S1/S2 cleaved S protein are visible at approx. 115 kDa.

### Residue E891 in the FP1 region seems critical for the fusogenic activity of MERS-CoV S

After we established that all mutated versions of the MERS-CoV S protein are expressed and exhibit a similar cleavage pattern as WT S, we initially screened whether the different mutant proteins are compromised in their ability to induce cell-cell fusion. Therefore, we co-expressed the S protein together with MERS-CoV receptor DPP4 in Vero cells for 18 h and visualized syncytia formation using an immunofluorescence assay (IFA). Trypsin treatment of transfected cells was not required as the cells natively express proteases that cleave WT and mutated S proteins for cell-cell fusion. As a control, we added the protease inhibitor dec-RVKR-CMK to cells expressing WT S to prevent S cleavage and block MERS-CoV entry [27]. We analyzed the cells by fluorescence microscopy and quantified cell-cell fusion by counting the number of nuclei per syncytium. Cells expressing WT S exhibited strong syncytia formation with multiple nuclei and addition of dec-RVKR-CMK as a control efficiently prevented cell-cell fusion (Figure 3A and B). Strikingly, in the absence of the inhibitor, the E891A and E891A/D892A mutants were impaired in their ability to trigger cell-cell fusion to the same extent as cells treated with dec-RVKR-CMK (Figure 3A and B). However, since the cells expressing D892A formed syncytia similar to that of cells expressing WT S, this suggests that E891 is a critical residue for the fusogenic activity of MERS-CoV S and was responsible for the E891A/D892A mutant’s inability to mediate cell-cell fusion. All other tested mutants showed attenuated cell-cell fusion at various degrees, however, none of the mutants was found to be as defective as the E891A S protein in mediating cell-cell fusion (Figure 3A and B).

**Figure 3:**
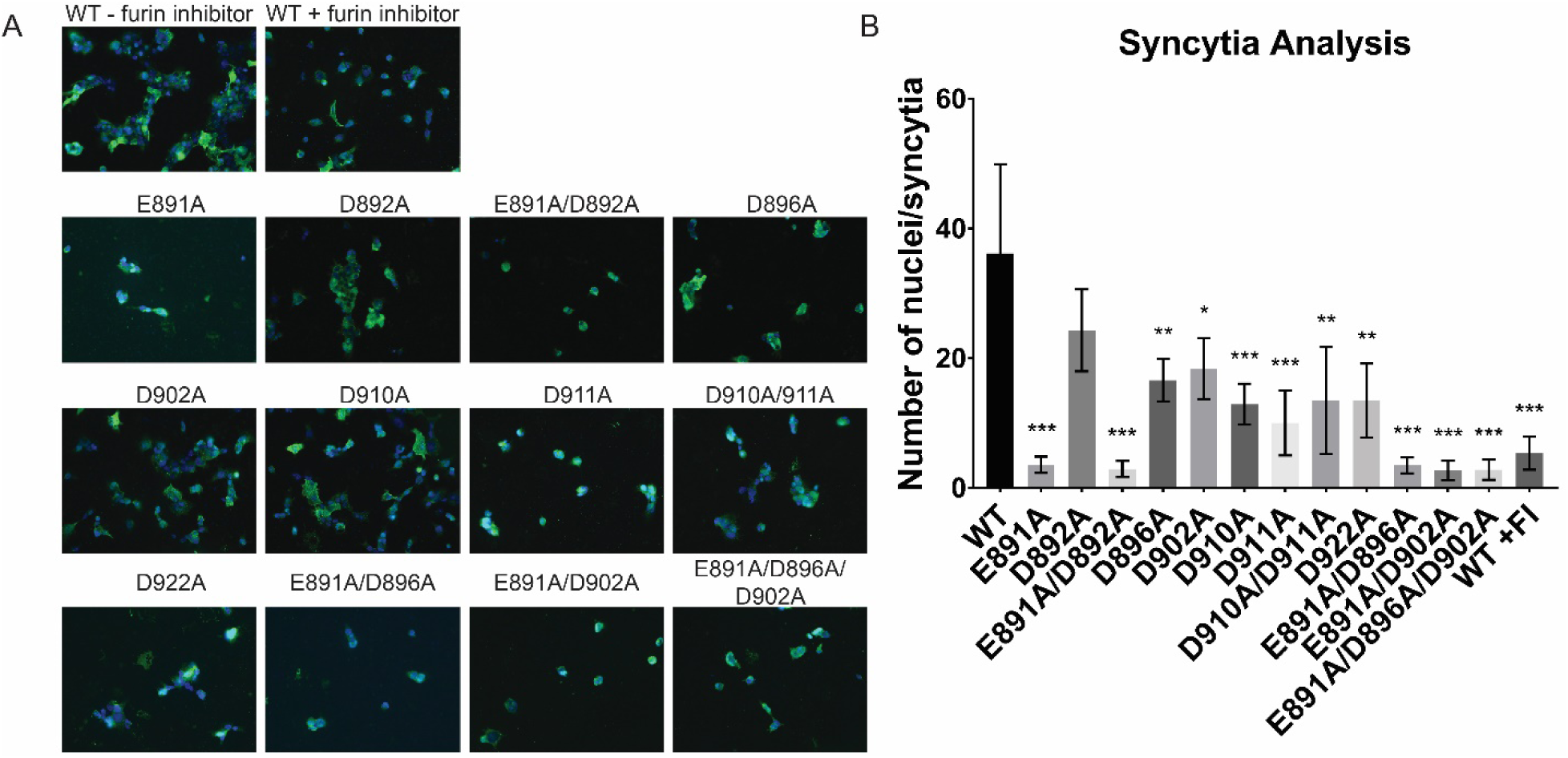
Immunofluorescence assay of MERS-CoV S WT and mutants. (A) Vero cells were transfected with plasmid DNA encoding for the respective MERS-CoV S variants and the DPP4 binding receptor and grown for 18 h. As Vero cells express endogenous proteases, which cleaves MERS-CoV S for fusion, no further protease treatment was needed to induce syncytia formation. WT + protease inhibitor indicates the condition in which protease inhibitor dec-RVKR-CMK at a concentration of 75 µM was added at the time of transfection to block fusion. Syncytia was visualized using immunofluorescence microscopy by staining the MERS-CoV S with a polyclonal anti-S antibody (in green) and the nuclei with 4′,6-diamidino-2-phenylindole (DAPI, in blue). Images were taken at a magnification of 25x. (B) Quantification of syncytia. Nuclei of 9 syncytia were counted and the average number of nuclei per syncytia was calculated. Error bars represent standard deviation (n = 9). Statistical analysis was performed using an unpaired student’s t-test comparing the WT against each of the respective mutant * = p > 0.5, ** = p > 0.05, *** = p > 0.005.

### Mutation of E891 diminishes infectivity of MERS-CoV pseudo particles

The cell-cell fusion results strongly suggest that E891 is an important residue for the fusogenic activity of MERS-CoV S. With the exception of D892, the data obtained for the other mutants indicate that they may play a minor role to trigger fusion. However, the mechanisms of cell-cell fusion and virus-cell fusion may significantly differ. Factors such as membrane curvature and/or density of viral S proteins may impact virus-cell fusion differently than cell-cell fusion [19]. Hence, the cell-cell fusion results provide an indication about the role of the individual D and E residues in the FP region, but need to be verified by viral infection of host cells. As MERS-CoV is a Risk Group 3 agent that needs to be handled in a biosafety level 3 (BSL-3) setting, we employed a viral pseudotyping technique to generate pseudo-particles (PP) that serve as surrogates of native virions and are suitable for infection in a BSL-2 setting. These pseudo-particles consist of a murine leukemia virus (MLV) core and are decorated with their respective S glycoprotein so that they recapitulate the entry steps (binding and fusion) of the native virus [28]. Furthermore, the PP carry a genome encoding for luciferase, and upon successful infection of cells, the luciferase reporter is integrated in the host cell genome and drives luciferase production within the cell, which can be used to quantify the degree of infectivity. We generated PPs carrying the mutant S proteins and infected Huh-7 cells, a permissive cell line for MERS-Cov infection with high DPP4 expression [15]. After 72 h, the cells were lysed and briefly incubated with the luciferin substrate, which can be oxidized with cellular luciferase to produce luminescence. The measured relative luminescence values provide a quantitative read-out for the infectivity of each mutant S protein mediated infection.

To ensure that the infectivity of PPs carrying different S protein variants can be compared to each other, we performed a Western blot analysis to detect the amount of S that was incorporated into the PPs (Figure 4). Because the cells were not treated with the protease inhibitor dec-RVKR-CMK, the S proteins were cleaved by endogenously expressed proteases. PPs with WT S and the mutant S proteins E891A, D892A, E891A/D892D, D896A, D902A, D910A, D911A, D910A/D911A and D922A exhibited comparable levels of incorporated full-length protein (∼ 250 kDa) and protein cleaved at the S1/S2 site (∼ 130 kDa). Cleavage at the S2’ site seemed slightly reduced for the D896A, D902A, D910A, D911A, D910A/D911A and D922A S protein mutants (∼ 115 kDa). As the Western blot data show that the PP incorporate similar amounts of full length and cleaved S proteins, for all S variants, we continued to analyze the infectivity of PPs carrying these mutated S proteins.

**Figure 4:**
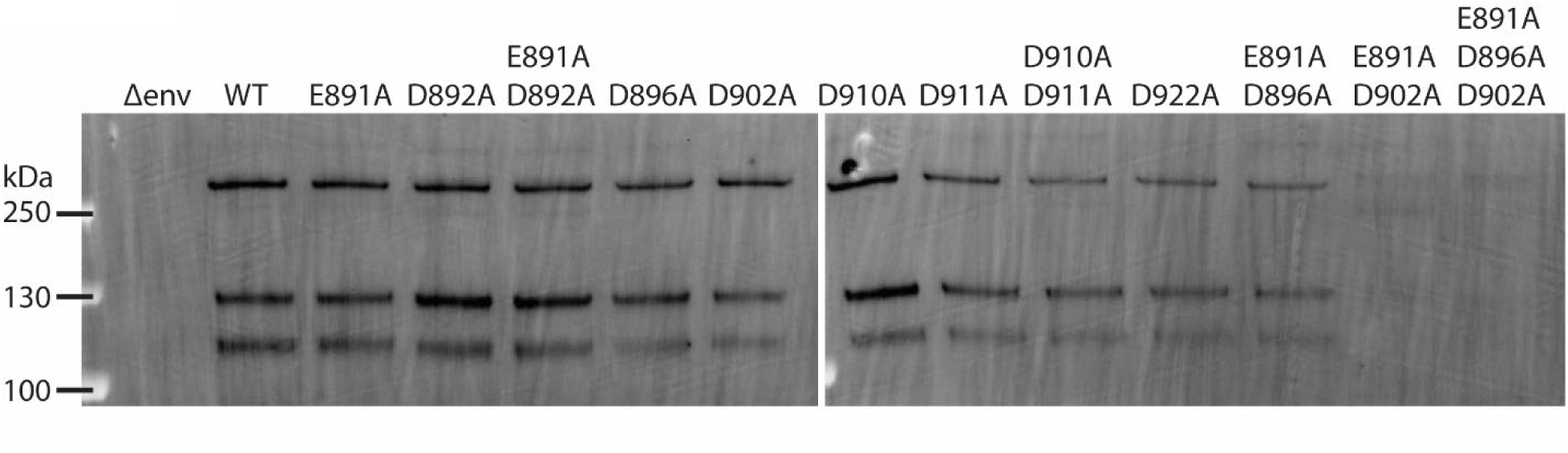
Western blot analysis of S proteins incorporated into PPs. 1 ml of DMEM containing PPs per each tested S protein were ultra-centrifuged, washed in PBS and resuspended in SDS Laemmli Buffer. Incorporated S proteins were analyzed using SDS-PAGE and detected using a Western blot with MERS-CoV S antibodies.

In agreement with the IFA observations, E891A and E891A/D892A PPs exhibited a five-fold reduction in the infectivity compared to the WT S PP, while D892A PP was not significantly different from WT S PP (Figure 5A). In addition, we also observed that infectivity of PPs carrying D911A was significantly lower than WT S PPs. However, infectivity was restored in the D911A/D912A double mutant (Figure 5A), suggesting that the D911 residue is not as critical for fusion as the D891 residue. If D911 was crucial for fusion, we would have expected the double mutant to display impaired infectivity as well. None of the other tested mutants exhibited significant differences compared to infections with WT S PPs (Figure 5A). D922A seemed to exhibit enhanced infection compared to the WT, but we did find it to be not statistically significant (Figure 5B).

**Figure 5:**
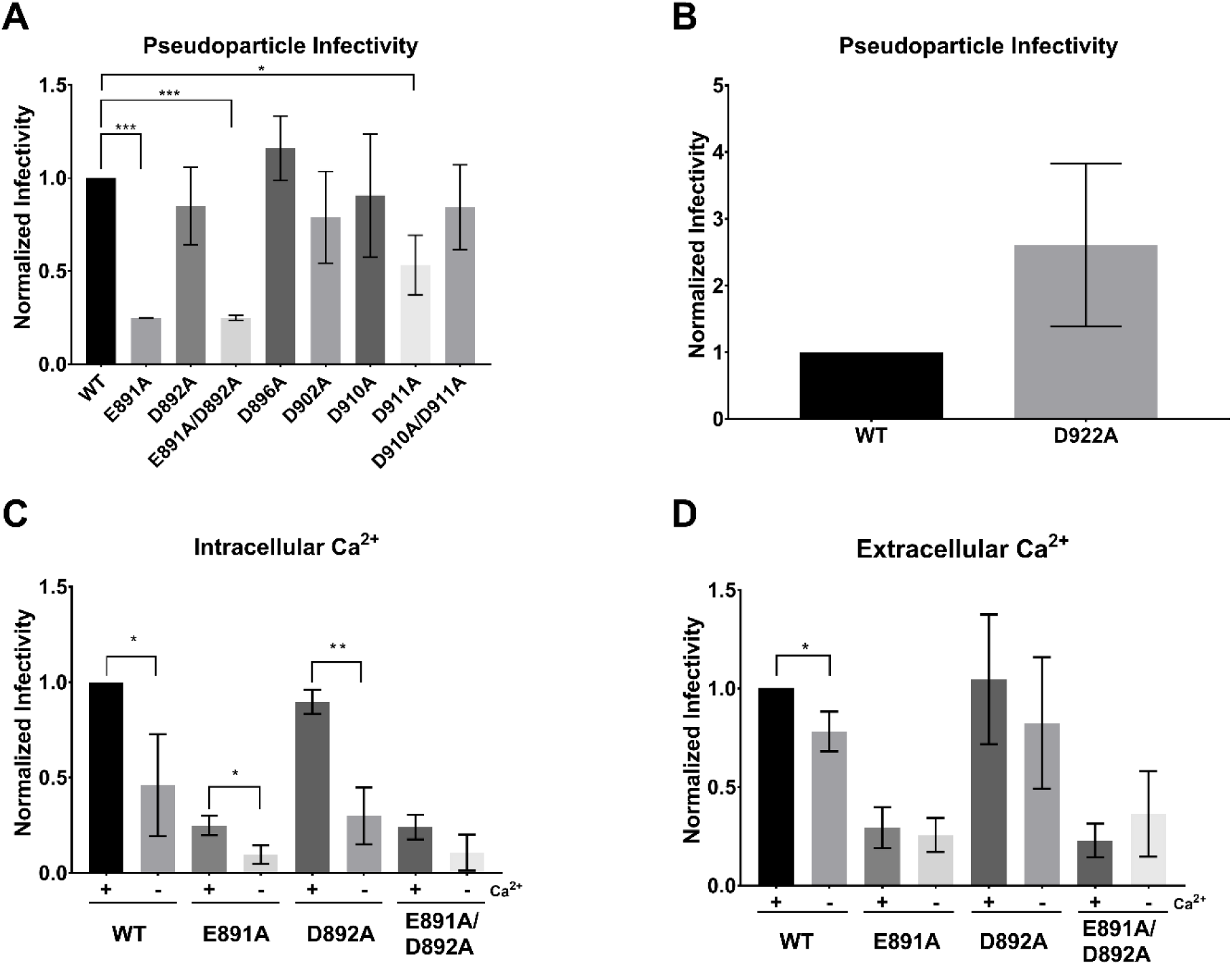
Pseudo-particle assays of MERS-CoV S WT and mutants. Huh-7 cells were infected with MLV-based pseudo-particles (PP) carrying MERS-CoV S WT or one of the respective S mutants. After 72 h, infected cells were lysed and assessed for luciferase activity. (A) PP infectivity of Huh 7 cells. (B) Infectivity of PP carrying the D922A S protein. Δenv and VSV-G served as representative controls for all PP assays (C) Impact of intracellular Ca^2+^ on MERS-CoV fusion. Cells were pre-treated with growth medium containing either 50 µM calcium chelator BAPTA-AM ordimethyl sulfoxide (DMSO) for 1 h. Cells were then infected with their respective PP in the presence of BAPTA-AM or DMSO for 2 h and grown for 72 h before assessing for luciferase activity. (D) Impact of extracellular Ca^2+^ on MERS-CoV fusion. Cells were pre-treated with growth medium either with or without 1.8 mM Ca^2+^ for 1 h. Infection protocol is as described above except PP were treated with 1.5 mM EGTA for calcium chelation. Infectivity was normalized such that WT PP infectivity is 1. Error bars represent standard deviation (n = 3). Statistical analysis was performed using an unpaired student’s t-test, as indicated. * = p > 0.5, ** = p > 0.05, *** = p > 0.005.

### Depletion of intracellular and extracellular Ca^2+^ ions abates viral infectivity

SARS-CoV was previously shown to depend on both extracellular and intracellular Ca^2+^ pools for successful infection, and it is believed that this dependency is due to Ca^2+^ interactions with the SARS-CoV FP [24]. Because of the high sequence similarity between the SARS-CoV and MERS-CoV FP, (Figure 1A) we were interested in determining if extracellular and/or intracellular Ca^2+^ also plays a role in MERS-CoV fusion. Our analysis of the negatively charged residues in the MERS-CoV S FP region thus far suggest that E891 is a crucial residue for viral fusion, perhaps for coordinating with Ca^2+^. We are further interested in observing if the fusion behavior of E891 mutants would differ from that of WT in varying calcium concentration environments. Thus, we focused only on WT S and S proteins carrying the amino acid substitutions E891A, D892A, and E891A/D892A. Although the D911A expressed decreased infectivity, we did not proceed for reasons previously mentioned.

First, we depleted intracellular Ca^2+^ by incubating the cells with 50 µM of 1,2-bis(2-aminophenoxy)ethane-*N,N,N’,N’,*-tetraacetic acid tetrakis (BAPTA-AM) prior to infection with PPs. BAPTA-AM is a Ca^2+^ chelator that is only active intracellularly and does not significantly impact cell viability at this concentration [24,29]. Chelating intracellular Ca^2+^ resulted in a two-fold drop in infectivity of MERS-CoV WT PPs (**Figure 5C**). Infectivity was also reduced in the E891A and the D892A mutants. However, we did not detect a significant reduction of infectivity in the E891A/D892A double mutant when depleting intracellular Ca^2+^. We next tested the impact of extracellular Ca^2+^ on MERS-CoV fusion with the host cell membrane by incubating the cells with 150 µM of ethylene glycol-bis(β-aminoethyl ether)-N,N,N’,N’-tetraacetic acid (EGTA) prior to infection. Infectivity of MERS-CoV WT PPs was reduced by 25% when extracellular Ca^2+^ was depleted (Figure 5D). There was no difference in infectivity with any of the PP carrying the mutated S proteins.

### E891A/D902A Mutation impairs S protein incorporation into PPs and subsequent infectivity

The results described above suggest that Ca^2+^ promotes entry of MERS-CoV into the host cell, with a greater dependency on intracellular Ca^2+^ concentration than extracellular Ca^2+^ concentration. Since we hypothesized that E891 is a critical residue in Ca^2+^ coordination, we would have expected its knockout variant, E891A, to have infectivity levels that are independent of Ca^2+^ concentrations. However, the intracellular Ca^2+^ studies show that single mutant E891A is impacted by Ca^2+^. A possible explanation for this apparent contradiction could be due to the FP flexibility. Perhaps if E891 is not available for Ca^2+^ binding, the adjacent D892 can compensate for E891 and maintain Ca^2+^ coordination in the FP. In support of this explanation is that the infectivity of the E891A/D892A double mutant is Ca^2+^ independent, as both possible residues that can mediate Ca^2+^ interactions have been knocked out. Thus, we believe that E891 does mediate Ca^2+^ binding in the MERS-CoV FP1 region. However, previous studies with RuV and EBOV showed that two distinct opposing residues in the FP region are required to coordinate Ca^2+^ [25,26], and so we proceeded to determine the other residues that Ca^2+^ can coordinate with.

MERS-CoV FP modeling suggests that there are two possible residues in the FP1 region that Ca^2+^ can coordinate together with E891, namely D896 and D902 (Figure 1B). Therefore, we created MERS-CoV S proteins carrying E891A/D896A, E891A/D902A, and E891A/D896A/D902A substitutions. We confirmed that all three mutated proteins are expressed and trafficked to the plasma membrane in 293T cells as proteins were extracted by cell-surface biotinylation (Figure 2C). The IFA results show that cells expressing E891A/D896A, E891A/D902A, and E891A/D896A/D902A are impaired in their ability to form syncytia to a similar extent as the cells expressing WT S treated with dec-RVKR-CMK or the E891A single mutant (Figure 3A and B). However, when we assessed protein incorporation into PPs, we found that WT and E891A/D896A PPs carried similar amounts of protein, but the protein incorporation in E891A/D902A and the E891A/D896A/D902A PPs was significantly reduced (Figure 4). When cells were infected with PPs equipped with the S mutations, we found that infectivity of E891A/D896A was similar to the E891A single mutant (Figure 6A). Infectivity of E891A/D902A, however, was significantly lower compared to E891A and E891A/D896A. PPs carrying the E891A/D896A/D902A S protein were not infectious. These findings are consistent with the results of the PP Western blot analysis since they incorporate significantly less S protein, which most likely results in reduced entry. Depleting intracellular Ca^2+^ levels using BAPTA-AM did not change infectivity of the tested PPs carrying S with the different amino acid substitutions (Figure 6B). When chelating extracellular Ca^2+^ using EGTA, infectivity was also not altered in the mutants (Figure 6C). In summary, our further mutant analysis to identify the second binding partner of Ca^2+^ are inconclusive. The data suggest that E896A is likely not the second binding partner because the E891A/D896A infectivity is similar to that of the E891A single mutant. However, due to poor incorporation of the E891A/D902A mutants into the PP, we are not able to compare their infectivity with the WT or E891A single mutants as those have higher amounts of S protein.

**Figure 6:**
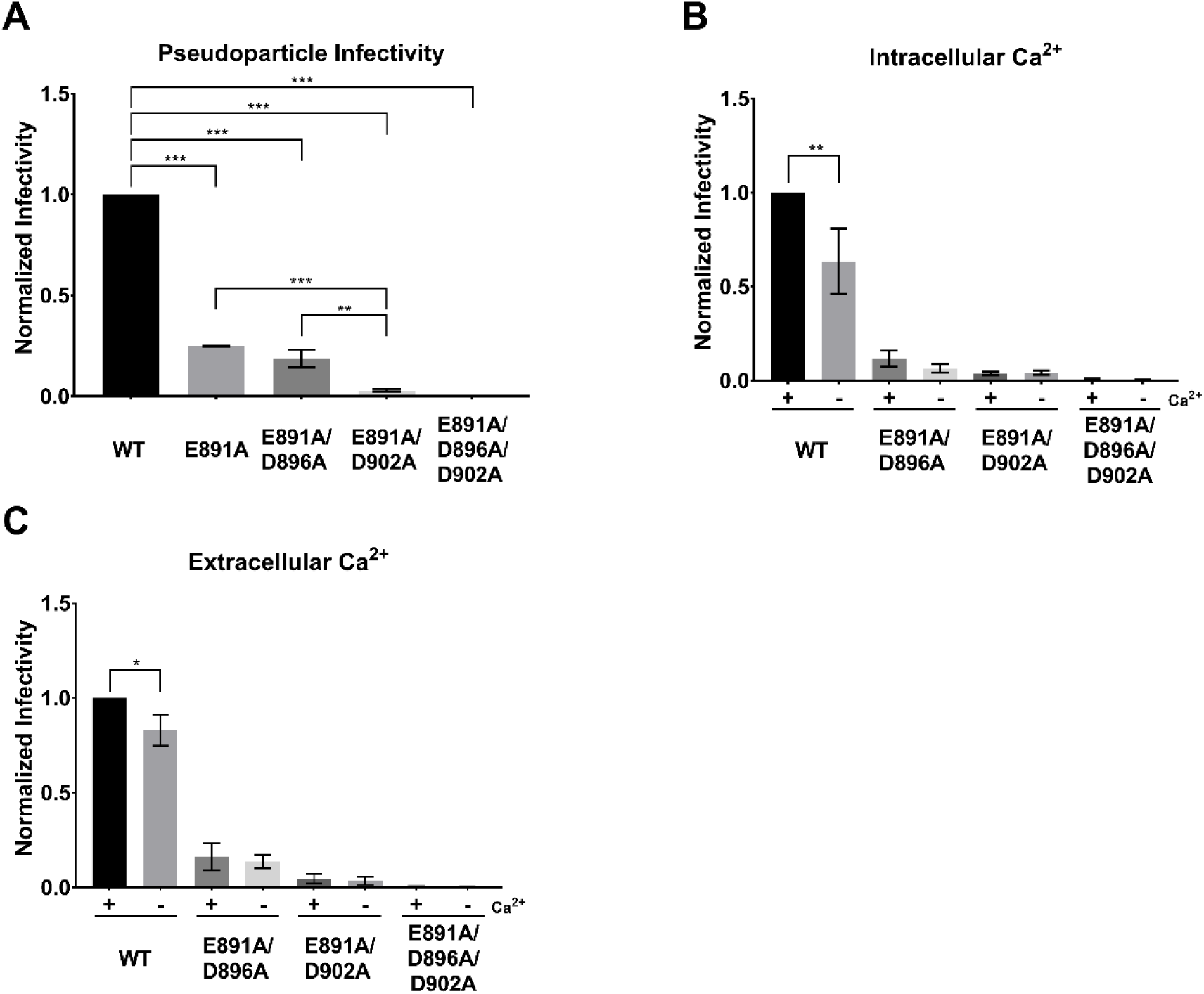
Pseudo-particle assays of MERS-CoV S WT and E891A/D896A, E891A/D902A and E891A/D896A/D902A mutants. Huh-7 cells were infected with MLV-based pseudo-particles (PP) carrying MERS-CoV S WT or one of the respective mutants. Infectivity was normalized to WT sample. Error bars represent standard deviation (n = 3). Statistical analysis was performed using an unpaired student’s t-test comparing the WT against the respective mutant (for B and C the untreated WT was compared to each sample). * = p > 0.5, ** = p > 0.05, *** = p > 0.005. (A) Infectivity of PPs without pre-treatment of cells. (B) Impact of intracellular Ca^2+^ on MERS-CoV fusion. Cells and PPs were treated as described for Figure 5 C. (C) Impact of extracellular Ca^2+^ on MERS-CoV fusion. Cells and PPs were treated as described for Figure 5 D.

### Ca^2+^ increases MERS-CoV FP membrane insertion and lipid ordering

We used electron spin resonance (ESR) to examine the effect of MERS-CoV FP on membranes and to test whether membrane fusion of the peptide requires Ca^2+^. In this technique, spin labels are incorporated into lipids at various positions, acting as depth probes. Multilamellar vesicles (MLVs) containing dipalmitoylphosphatidyl-tempo-choline (DPPTC) are spin labeled in the head region whereas those with phosphocholine are labeled in either the upper tail region (5PC) or lower tail region (14PC). The ordering parameter, S_0_, is an indication of the amount of membrane ordering at a given depth and serves as a readout for FP membrane penetration depth [30–32]. Changes in membrane ordering may reduce the energy barrier between two membranes and thus promote fusion [31,33]. Previously studied FPs from different viruses induced membrane ordering illustrated as an S-shaped curve of S_0_ as a function of increasing FP to lipid (P/L) ratio while non-functional FPs and random peptides do not have a membrane-ordering effect and produce a scrambled signal [24,30–32,34]. Therefore, examining the membrane ordering effect of a peptide allows to determine whether it is a FP [24,34].

As shown in Figure 7A and B, MERS-CoV FP induced the greatest ordering effect of the head (DPPTC) and upper-tail (5PC) regions with 1 mM Ca^2+^. When we tested the deep hydrophobic region (14PC), no membrane ordering was observed (data not shown). When no Ca^2+^ is supplemented and trace Ca^2+^ is chelated with 1 mM EGTA at the same time, no such ordering effect is measured. This indicates that the function of the MERS-CoV FP is Ca^2+^ dependent. Compared to the SARS-CoV FP, however, the MERS-CoV FP has a weaker effect. For DPPTC case, the maximal S_0_ for MERS-CoV FP is smaller (approx. 0.472) than that for SARS-CoV FP (approx. 0.479) at the highest P/L ratio in our experiment (5%). We fitted the S-shape curve using a four parameter logistic equation Y=S_0,min_+(S_0,max_-S_0,min_)/(1+(X-X_0_)^p^), which is a built-in function of Origin (OriginLab), where the X_0_ is the P/L ratio (X) that induces half of maximal membrane order change (S_0,max_-S_0,min_), and p is the Hill’s slope. The Hill’s slope is an indicator for cooperativity in a ligand-receptor binding reaction. The X_0_ for SARS-CoV FP is 0.25% and for MERS-CoV FP is 0.61%, indicating the MERS-CoV FP is less effective than the SARS-CoV FP. The Hill’s slope for SARS-CoV FP is 2.39 and for MERS-CoV FP is 1.44, indicating SARS-CoV FP induces membrane ordering in a more cooperative fashion then MERS-CoV FP at 1 mM Ca^2+^. The scrambled peptide shows no effect on membrane order. Although the scrambled control peptide is based on the SARS-CoV FP sequence, it is used as a control for MERS-CoV FP, too, because of the high sequence similarity of MERS-CoV and SARS-CoV FPs (Figure 1A).

**Figure 7:**
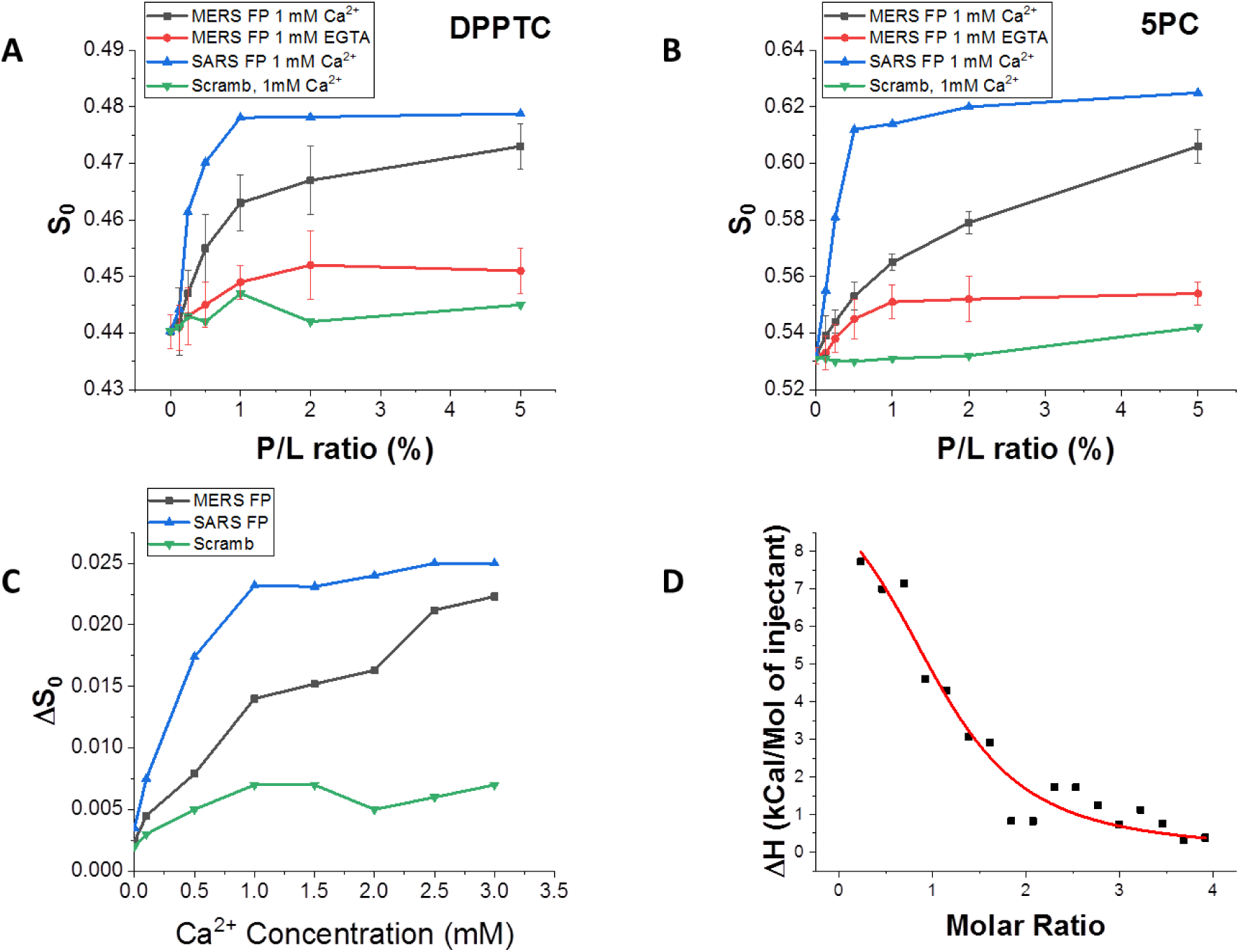
ESR and ITC analysis of the MERS-CoV FP. A-B. Plots of order parameters of DPPTC (A), and 5PC (B) versus peptide:lipid ratio (P/L ratio) of MERS FP or SARS FP in POPC/POPS/Chol=3/1/1 MLVs in buffer with 150 mM NaCl at 25°C. Black, MERS FP, 1 mM Ca^2+^ and at pH 5; red, MERS FP calcium-less buffer with 1 mM EGTA and at pH 5; blue, SARS FP, 1 mM Ca^2+^ at pH 5, and purple, scrambled peptide, 1 mM Ca^2+^ and at pH 5. (C) Plot of difference of order parameters of DPPTC with and without 1% peptide binding (ΔS0) versus Ca^2+^ concentration in POPC/POPS/Chol=3/1/1 MLVs in buffer with 150 mM NaCl at 25°C. Black, MERS FP; blue, SARS FP; and green, scrambled peptide. The experiments were typically repeated two to three times. The typical uncertainties found for S_0_ ranges from 1-5 × 10^−3^, while the uncertainties from repeated experiments were 5-8 × 10^−3^ or less than ±0.01. We show the standard deviation bars in Panel A and B. (D) ITC analysis of Ca^2+^ binding to MERS-CoV FP. The peptides were titrated with CaCl_2_. The integrated data represent the enthalpy change per mole of injectant, ΔH, in units of kJ/mol as a function of the molar ratio. Data points and fitted data are overlaid. The fitting is based on the one-site model.

To further test the effect of Ca^2+^ on S_0_, we varied the Ca^2+^ concentration while fixing the P/L ratio at 1%. The effects of any ordering caused by Ca^2+^ interacting with the liposomes rather than the fusion peptide was removed by subtracting S_0_ of a liposome and Ca^2+^ only (no peptide) mixture from the measurement of S_0_ when all three components (liposomes, peptides, and Ca^2+^) were present, yielding ΔS_0_. This method proved to be a solid approach to single out the effect of FP under the influence of Ca^2+^ in a complex membrane-peptide-Ca^2+^ system [24,26]. As shown in Fig. 7C, ΔS_0_ increased with increasing Ca^2+^, indicating that there is more FP-induced membrane ordering at higher Ca^2+^ concentrations in the lipid head group regions. When fitted with the same logistic equation, we calculated the X_0,Ca_ (the concentration of Ca^2+^ to enable half of the maximal FP-induced ordering effect) for SARS-CoV is 0.32 mM. Although the X_0,Ca_ for MERS-CoV has a very large standard error and thus is not reliable, due to the fact that the ΔS_0_-[Ca^2+^] curve has not been saturated within our experimental range (0-3 mM). But X_0,Ca_ for MERS-CoV is obviously larger than that for SARS-CoV. The values for SARS-CoV FP and MERS-CoV FP are 1.56 and 0.70, respectively, indicating that Ca^2+^ ions bind to the SARS-CoV FP more cooperatively than to MERS-CoV FP. For the SARS-CoV FP case, an explanation for this cooperativity is that the SARS-CoV FP has two Ca^2+^ binding sites, and binding a Ca^2+^ ion in one site will stabilize the secondary structure and help to bind another Ca^2+^ ion in the other site. The lack of cooperativity in the case of the MERS-CoV FP suggests that it may have only one binding site.

### The binding stoichiometry of Ca^2+^ and MERS CoV FP is close to one

To directly investigate the interaction between Ca^2+^ and MERS-CoV FP, we used isothermal titration calorimetry (ITC). In this experiment, the synthetic MERS-CoV FP was added into the reaction cell at neutral buffer conditions (pH 7) and 1 ml CaCl_2_ (pH 7) was titrated into the cell dropwise. The heat flows generated by the reaction were recorded and the reaction heat for each injection (ΔH) is calculated by integrating the heat flow over the time. After the subtraction of dilution heat, which is measured by titrating 1 ml CaCl_2_ into the pH 7 buffer, the reaction heat between the Ca^2+^ and the FP for each injection vs. the molar ratio of Ca^2+^ to FP was plotted. As shown in Figure 7D, the reaction heats are positive, indicating the reaction is endothermic. The reaction heats decrease during the titration, indicating the FPs have been saturated by the Ca^2+^ ions toward the end of the titration. The reason for the endothermic reaction may be due to the Na^+^ ions interacting with the FPs were replaced by the Ca^2+^ ions. The endothermic reactions were also observed when we injected CaCl_2_ into the SARS CoV FP in a similar experimental setup [24]. Fitting the data with the one-site model, which assumes all binding site(s) have the same binding affinity, we calculated the binding constant K_b_=1.14±0.23 ×104 M^−1^, the reaction enthalpy ΔH= 3.12 ± 0.45 kCal/Mol and the binding stoichiometry (Ca^2+^ to the FP) n=1.12±0.08. The stoichiometry of the Ca^2+^ to the MERS-CoV FP is significantly different from that for the FP1_2 of SARS-CoV FP (n=1.7) and similar to FP1 or FP2 of the SARS-CoV FP (n=0.92 and 1.02, respectively) [24]. The results strongly support the hypothesis that the MERS-CoV FP only binds one Ca^2+^ ion, while the SARS-CoV FP binds two, one on FP1 and one on FP2. The weaker Ca^2+^ binding energies of the MERS-CoV FP is also consistent with the ESR results, which shows that the MERS-CoV FP induces a maximal membrane ordering effect in a higher Ca^2+^ concentration than the SARS-CoV FP.

## Discussion

Viral entry into the host cell is one of the most critical steps in the life cycle of a virus. Enveloped viruses such as MERS-CoV and SARS-CoV utilize class I fusion proteins that facilitate binding to and fusion with the host cell membrane. A central factor that allows the viral and host membrane to merge is the FP within these proteins that is exposed upon proteolytic cleavage. Recently, reports demonstrated a crucial role for Ca^2+^ in viral fusion, as it coordinates with negatively charged residues such as D and E within the FP. Here, we show that Ca^2+^ promotes viral entry of MERS-CoV into the host cell. Of the investigated residues, E891 seems to be the major residue for Ca^2+^ binding because the E891A substitution dramatically decreased viral infectivity (Figure 5A). We believe, however, due to the flexibility of the FP that the adjacent D892 residue can compensate for the loss of E891 because we observed attenuated infectivity in E891A when depleting intracellular Ca^2+^ levels that was restored in the E891A/D892A mutant (Figure 5C). The shorter side chain of D at the 892 position may not establish a bond with Ca^2+^ as strong as E and therefore the restoration of infectivity is only very minimal.

Our attempts to identify the binding partner by generating double mutants of E891 with neighboring residues D896 and D902 were not successful as we observed poor S protein incorporation into the PPs. It is intriguing that the Western blots and cell-cell fusion results show that the E891A/D902A S mutants are properly expressed and trafficked to the cell membrane but are poorly integrated the virion surrogates. One possible explanation is that these mutations impacts the integrity of the FP domain in a manner that hinders virion incorporation.

The role of Ca^2+^ for viral fusion has been explored for other enveloped viruses such as RuV [25], SARS-CoV [24], EBOV [26], human immunodeficiency virus type I (HIV-1) [35] and influenza A [24]. There seem to be significant differences regarding the role of Ca^2+^ in mediating viral host cell membrane fusion. Studies showed that Ca^2+^ does not bind to the FPs of influenza A or HIV-1 during the fusion event. Our prior observations of influenza fusion protein, which does not show increased membrane interaction in the presence of Ca^2+^ [24], indicate that Ca^2+^ does not act as a blanket promoter of viral FP insertion into target membranes. [24]. In the case of HIV this leads to structural changes of the FP, which forms an α-helix in the absence of Ca^2+^, but changes conformation to an antiparallel β-sheet structure in the presence of Ca^2+^ [35]. However, it is important to note, that while Ca^2+^ enhances fusion for the above-mentioned viruses it is attenuated in its absence, but not abrogated. In contrast, Ca^2+^ directly interacts with the FPs of RuV, SARS-CoV, and EBOV and depletion of Ca^2+^ results in the incapability of the virus to fuse with the host cell [24–26]. For both RuV and EBOV, key residues within the FP were identified that coordinate with the Ca^2+^, and in concert with low pH, resulted in conformational changes of the FP and subsequent fusion. Here, we demonstrate that MERS-CoV FP has a similar interaction with Ca^2+^ as SARS-CoV FP but with distinct differences.

Our data shows that Ca^2+^ significantly promotes MERS-CoV fusion and that fusion in the absence of Ca^2+^ is attenuated, but not completely abrogated. This could be due to the fact that the MERS-CoV FP binds only one Ca^2+^ ion (which is discussed in greater detail below). On the opposite, host cell entry of SARS-CoV which binds two Ca^2+^ in its FP region is entirely abrogated when depleting intracellular Ca^2+^ levels [24]. Given the high sequence similarity between the FP regions of the two viruses this finding implicates that the stoichiometry between FP and Ca^2+^ has a significant impact on the fusion dynamics of the virus. However, because MERS-CoV shows a similar fusion behavior as influenza A or HIV-1 in the absence of Ca^2+^, it could be argued that fusion of MERS-CoV is regulated by Ca^2+^ to the membrane. We believe that we can rule out this possibility because our data obtained with the E891A mutant, the ESR and ITC provide good evidence that Ca^2+^ interacts directly with the MERS-CoV FP.

By using ESR, we have demonstrated that the MERS-CoV FP also exerts a membrane ordering effect similar to that observed for SARS-CoV FP as well as for other viral FPs, which further strengthens our hypothesis that the membrane ordering effect is a required step for viral entry [24]. The membrane ordering effect is in turn serves as a useful biophysical indicator for examining the conditions required for the FP activity. Using this method, we have previously demonstrated that SARS-CoV and EBOV FPs are Ca^2+^ dependent, and now show the MERS-CoV FP is Ca^2+^ dependent too. However, although the S-shaped curve of the MERS-CoV FP is similar to the SARS-CoV FP, MERS-CoV FP has a smaller membrane order effect than SARS-CoV FP and at 1.0 mM Ca^2+^, the membrane ordering effect is saturated by a higher P/L ratio of MERS-CoV FP than SARS-CoV FP (Figure 7A and 7B). By keeping the P/L ratio constant at 1%, the MERS-CoV FP is also saturated at a higher concentration of Ca^2+^ (increase to half of its maximal at around 1.5 mM, compared to around 0.5 mM for SARS-CoV FP, Figure 7C). Those results suggest that the MERS-CoV FP has a weaker interaction with Ca^2+^ and thus require higher Ca^2+^ concentration reach its maximal activity. This hypothesis is further supported by the ITC experiment, which shows that the reaction energies between the Ca^2+^ and the MERS-CoV FP are weaker than those of the SARS-CoV FP. The ITC experiments also demonstrate that the stoichiometry of Ca^2+^ to the MERS-CoV FP is close to one (1.15). This is different to the SARS-CoV case, in which SARS-CoV binds two Ca^2+^ in its FP region, one in each FP1 and FP2 [24]. We interpret this to indicate that one Ca^2+^ interacts with the MERS-CoV FP.

Thus, although the sequences of SARS and MERS-CoV FP are similar, they may interact with Ca^2+^ differently. We believe that this interaction occurs in the MERS-CoV FP1 region as our data identified a residue in the FP1 region that interacts with Ca^2+^ suggesting that this domain is important for Ca^2+^ binding. None of the tested mutants in the FP2 region provided clear evidence that Ca^2+^ binding occurs in this domain, which is different from SARS-CoV FP. It is striking that despite the high sequence similarity between SARS-CoV and MERS-CoV FP, there are significant differences in their FP Ca^2+^ interactions. The ESR results indicate that the ordering effect is sequence specific. It is possible that the small difference in the sequence of the FP2 region of MERS-CoV and SARS-CoV is responsible for this difference in Ca^2+^ interaction.

We show here that while SARS-CoV and MERS-CoV both functionally interact with Ca^2+^ ions, there appear to be significant positional and stoichiometric differences between the two spike proteins. Prior mutagenesis experiments were carried out on the conserved E and D residues in the SARS-CoV FP1-2 region before the recognition of the bipartite nature of the FP and prior to any consideration of their potential for Ca^2+^ interaction [36,37]. For SARS-CoV, the residue equivalent to MERS-CoV E891 (E801) could not be incorporated into PPs when mutated. Instead, the FP1 SARS-CoV residue D812 showed a major reduction in fusion activity when mutated. The equivalent MERS-CoV residue (D902) only showed a much more modest effect when mutated. Notably, SARS-CoV D830 (in what is now defined as FP2) also showed a major reduction in fusion activity when mutated, in contrast to the equivalent residue in MERS-CoV (D922). D922A actually showed increased infectivity when mutated, the reasons for this are currently unclear. Overall, these prior studies are in broad agreement with the interaction of Ca^2+^ with only FP1 for MERS-CoV, compared to both FP1 and FP2 for SARS-CoV.

In the biological scenario, a possible explanation for the differences might be that Ca^2+^ primes the FP for different steps in the host cell entry process. For SARS-CoV, extracellular Ca^2+^ binding through the FP2 domain may prepare the fusion loop for subsequent intracellular Ca^2+^ in the FP1. For MERS-CoV, the extracellular Ca^2+^ binding step may not be required as none of the tested mutants in the FP2 region showed a striking reduction on infectivity whereas intracellular Ca^2+^ depletion significantly reduced infectivity. Future studies will be designed to explore how swapping different residues or entire regions between MERS-CoV and SARS-CoV FP would impact membrane fusion, entry pathways, and calcium interactions. It would be interesting to see if MERS-CoV will adopt SARS-CoV fusion behavior and vice versa. Different cell types, however, could also affect the fusion behavior of both viruses. Previous studies explored the fusion dynamics of SARS-CoV and MERS-CoV PP using Vero E6 cells [24,38] while we and others used Huh-7 cells to study MERS-CoV PP infections which exhibit a higher expression of furin and DPP4 and seem to promote entry mainly via the endocytic route [16,39]. Addressing the impact of the cell type on the fusion dynamics of MERS-CoV and SARS-CoV will complement our studies on the FP domains of the two viruses.

## Acknowledgements

This work was supported by the National of Health research grant R01AI35270, R01GM123779, and P41GM103521. We also would like to thank Dr. David Eliezer (Weill Cornell Medicine) and Javier A. Jaimes (College for Veterinary Medicine, Cornell) for their critical input. We thank Dr. Brian Crane (Cornell University, Department of Chemistry and Chemical Biology) for sharing equipment.

## Materials and Methods

### Cells, plasmids, and reagents

HEK293T and Vero-E6 cells were obtained from the American Type Culture Collection. Huh-7 cells were obtained from the Japan Health Science Research Resources Bank. All cells were maintained in Dulbecco’s modified Eagle medium (DMEM) (Cellgro) supplemented with 25 mM HEPES (Cellgro) and 10% HyClone FetalClone II (GE) (cDMEM) at 37°C and 5% CO_2_. The MERS-CoV S EMC/2012 gene was codon-optimized for mammalian expression and cloned into pDNA3.1+ (Invitrogen) as described previously [39]. The pCMV-MLV gag-pol murine leukemia virus (MLV) packaging construct, the pTG-Luc transfer vector encoding luciferase reporter and pCAGGS/VSV-G plasmid were described before [15,40].

Recombinant L-1-Tosylamide-2-phenylethyl chloromethyl ketone (TPCK)-treated trypsin was purchased from Sigma. The protease inhibitor dec-RVKR-CMK was obtained from Tocris. Calcium chelators BAPTA-AM and EGTA were purchased from Tocris and VWR, respectively.

### Site-directed mutagenesis

All mutated MERS-CoV S constructs were generated by site directed mutagenesis using the QuikChange Lightning site-directed mutagenesis kit (Agilent). pDNA3.1+/MERS-CoV S EMC/2012 served as template for all the mutagenesis performed. PCR reactions and *E. coli* XL 10 Gold transformations were carried out according to the manufacturer’s recommendations. Primers (IDT) were designed using the primer design tool from Agilent. The following primer pairs were used: E891A (ATCGAACAACAGATCCGCGATCGCTGATCGTGC/GCACGATCAGCGATCGCGGATCTGTTGTTCGAT); D892A (CCTTATCGAACAACAGAGCCTCGATCGCTGATCGT/ACGATCAGCGATCGAGGCTCTGTTGTTCGATAAGG); E891A/D892A (CCTTATCGAACAACAGAGCCGCGATCGCTGATCGTGCG/CGCACGATCAGCGATCGCGGCTCTGTTGTTCGATAAGG); D896A (CGGCAATGGTCACCTTAGCGAACAACAGATCCTCG/CGAGGATCTGTTGTTCGCTAAGGTGACCATTGCCG); D902A (GCATATAGCCCGGAGCGGCAATGGTCACC/GGTGACCATTGCCGCTCCGGGCTATATGC); D910A (CCTGCTGCATGCAGTCAGCGTAACCTTGCATATAG/CTATATGCAAGGTTACGCTGACTGCATGCAGCAGG); D911A (CCCTGCTGCATGCAGGCATCGTAACCTTGCA/TGCAAGGTTACGATGCCTGCATGCAGCAGGG); D910A/D911A (GCCCCTGCTGCATGCAGGCAGCGTAACCTTGCATATAG/CTATATGCAAGGTTACGCTGCCTGCATGCAGCAGGGGC); D922A (GGCGCAGATAAGGGCTCTGGCGCTGGC/GCCAGCGCCAGAGCCCTTATCTGCGCC); E891A/D896A (GGCAATGGTCACCTTAGCGAACAACAGATCCGC/GCGGATCTGTTGTTCGCTAAGGTGACCATTGCC); E891A/D902A (GCATATAGCCCGGAGCGGCAATGGTCACC/GGTGACCATTGCCGCTCCGGGCTATATGC); E891A/D896A/D902A (GCATATAGCCCGGAGCGGCAATGGTCACC/GCGGATCTGTTGTTCGCTAAGGTGACCATTGCC/GGCAATGGTCACCTTAGCGAACAACAGATCCGC/GGTGACCATTGCCGCTCCGGGCTATATGC). All constructs were verified by Sanger sequencing.

### Western blot analysis of MERS-CoV S WT and mutants

HEK293T cells were grown in 6-well plates and transfected with pCDNA3.1+ encoding the respective S protein variant using polyethylenimine (PEI) (Fisher). For each well, 2 µg DNA and 6 µL PEI were incubated with 50 µL Opti-MEM (Gibco) seperately for 5 minutes before combining and incubating for 20 minutes. After 20 minutes, 2 mL cDMEM was added to the transfection mixture. Cells were washed with phosphate-buffered saline (PBS) (Cellgro) once and the DNA/PEI/DMEM mix was added. For cells treated with protease inhibitor, 75 µM dec-RVKR-CMK was added to cells at the time of transfection. Cells were then incubated for 18 h. For cells treated with TPCK-trypsin, they were washed with PBS once, and 1 mL PBS supplemented with 0.8 nM TPCK-trypsin was added to the cells. Cells were then incubated for 10 min at 37°C and 5% CO_2_. Cell-surface biotinylation, western blotting, and antibody analysis were performed as previously described [41]. S protein was detected using the MERS-CoV S rabbit polyclonal antibody (Sino Biological, Cat No: 40069-RP01) as the primary antibody, and AlexaFluor 488-labeled anti-rabbit secondary antibody (Invitrogen).

### Immunofluorescence assay of MERS-CoV S WT and mutants

Vero cells were grown in microscopy chamber slides (Millipore) and transfected with the respective pCDNA3.1+/MERS-CoV S construct using Lipofectamine 2000 (Invitrogen). Transfections were performed according to the manufacturer’s protocol. 18 h post transfection, the immunofluorescence assay was carried out as previously described with the exception that membranes were not permeabilized with TritonX-100 (Millet PNAS). The MERS-CoV S protein was detected using the same antibodies as described for the western blot analysis. For quantification, images of at least five randomly selected fields were acquire. The nuclei of nine syncytia for each condition were counted manually to calculate the average nuclei/syncytia. Average and standard deviation were calculated using Microsoft Excel and GraphPad Prism 7. Data was visualized using GraphPad Prism 7.

### Pseudo-particle assays

Production of and infection with pseudo-particles (PPs) was performed as previously described with minor modifications [28]. In brief, PPs of each individual construct were produced by transfecting HEK293T cells with 600 µg of the respective plasmid DNA, 600 µg pTG-Luc and 800 µg pCMV-MLVgag-pol using PEI as described above. 48 h post transfection, the supernatants were harvested, centrifuged at 290 x g for 7 minutes to remove cell debris, filtered through a 0.45 µm syringe filter and stored at -80°C in small aliquots. For infection assays, Huh-7 cells were seeded in 24-well plates. Cells were washed with 500 µL PBS and 200 µL PPs were added and incubated for 2 h at 37°C and 5% CO_2_ with rocking. After 2 h, the cells were supplemented with 300 µL cDMEM medium and incubated for 72 h at 37°C and 5% CO_2_. Cells were then lysed, and luciferase activity was measured using a Luciferase Assay Kit (Promega). Readings were performed with a Glomax 20/20 system (Promega). For each experiment, three technical replicates were prepared. Each experiment was repeated at least three times. Data was analyzed using GraphPad Prism 7.

To chelate intracellular calcium, cells were washed with 500 µL PBS and subsequently pre-treated with 200 uL of 50 µM BAPTA-AM (dissolved in DMSO) in DMEM+ (2% Hyclone, 10 mM HEPES, 1.8 mM Ca^2+^) for 1 h at 37°C and 5% CO_2_. After 1 h, DMEM+ was removed and cells were infected with 200 µL PP containing 50 µM BAPTA-AM and incubated for 2 h at 37°C and 5% CO_2_ with rocking. After 2 h, cells were supplemented with 300 µL cDMEM and incubated for 72 h. Analysis of infected cells are as described above. Negative controls were conducted by pre-treating in DMEM+ and infecting with PP with an equivalent volume of DMSO.

To chelate extracellular calcium, cells were pre-incubated with 200 uL of DMEM- (2% Hyclone, 10 mM HEPES) for 1 h at 37°C and 5% CO_2_. After 1 h, DMEM-was aspirated and cells were infected with 200 µL PP containing 50 uM EGTA for 2 h 37°C and 5% CO_2_with rocking. Analysis of infected cells is as described above. Negative controls were controls were conducted by pre-incubating the cells in DMEM+ and infecting with PP without treatment. For Western Blot analysis of the incorporated S protein 1 ml of harvested PP were spun down at 43000 x g for 2 h at 4°C. The supernatant was aspirated, and protein pellet was resuspended in 2x Laemmli buffer. All protein was loaded on a SDS-PAGE gel and Western blot analysis was carried out as described above.

### Modelling of MERS-CoV S monomer

The structure of the MERS-CoV S protein was downloaded from the Protein Data Bank (PDB) (Accession Number 5X5C, DOI: http://dx.doi.org/10.2210/pdb5X5C/pdb) [42,43]. Modelling was performed as described previously using Chimera software (Chimera version 1.14, University of California) [44,45]. Image was created using Adobe Illustrator CC 22.1.

### Lipids and Peptides

The lipids POPC, POPS, and the chain spin labels 5PC, 10PC and 14PC and a head group spin label dipalmitoylphospatidyl-tempo-choline (DPPTC) were purchased from Avanti Polar Lipids (Alabaster, AL) or synthesized by our laboratory according to previous protocols. Cholesterol was purchased from Sigma (St. Louis, MO). All peptides were synthesized by SynBioSci Co. (Livermore, CA). The Sequence of SARS FP and its corresponding shuffle peptide are the same as in [24].

### Vesicle Preparation

The composition of membranes used in this study is consistent with our previous study [24]. The desired amount of POPC, POPS, cholesterol and 0.5% (mol:mol) spin-labeled lipids in chloroform were mixed well and dried by N_2_ flow. The mixture was evacuated in a vacuum drier overnight to remove any trace of chloroform. To prepare MLVs, the lipids were resuspended and fully hydrated using 1 mL of pH 7 or pH 5 buffer (5 mM HEPES, 10 mM MES, 150 mM NaCl, and 0.1 mM EDTA, pH 7 or pH 5) at room temperature (RT) for 2 hours. To prepare SUVs, the lipids were resuspended in pH 7 or pH 5 buffer and sonicated in an ice bath for 20 minutes or when the suspension became clear. The SUVs solution was then further clarified by ultracentrifugation at 13,000 rpm for 10 min.

### Isothermal Titration Calorimetry (ITC)

ITC experiments were performed in an N-ITC III calorimeter (TA Instrument, New Castle, DE). A total 97 µL of 1 mM CaCl_2_ in pH 7 fusion buffer was injected into 1 ml 0.05 mM FPs in pH 7 fusion buffer at 37°C in a stepwise manner consisting of 5 µL per injection except that the first injection was 2 μL. The injection time was 15 sec for each injection and the interval time was 10 min. The data were analyzed with Origin (OriginLab Corp., Northampton, MA). The one-site model was used in the fitting to calculate the thermodynamic parameters. The protein concentration is determined by dry weight and UV spectroscopy.

### ESR spectroscopy and nonlinear least-squares fit of ESR spectra

To prepare the samples for lipid ESR study, the desired amounts of FPs (1 mg/mL) were added into the lipid MLVs dispersion. After 20 min of incubation, the dispersion was spun at 13,000 rpm for 10 min. The concentrations of peptide were measured using UV to ensure complete binding of peptide. The pellet was transferred to a quartz capillary tube for ESR measurement. ESR spectra were collected on an ELEXSYS ESR spectrometer (Bruker Instruments, Billerica, MA) at X-band (9.5 GHz) at 25°C using a N_2_ Temperature Controller (Bruker Instruments, Billerica, MA).

The ESR spectra from the labeled lipids were analyzed using the NLLS fitting program based on the stochastic Liouville equation using the MOMD or Microscopic Order Macroscopic Disorder model as in previous studies [31]. The fitting strategy is described below. We employed the Budil *et al.* NLLS fitting program [46] to obtain convergence to optimum parameters. The g-tensor and A-tensor parameters used in the simulations were determined from rigid limit spectra [24]. The ordering tensor parameters, S_0_ and S_2_, were extracted from the spectra, which are defined as follows: S_0_=<D_2,00_>=<1/2(3cos2θ-1)>, and S_2_=<D_2,02_+D_2,0-2_> =<√(3/2)sin2θcos2f>, where D_2,00_, D_2,02_, and D_2,0-2_ are the Wigner rotation matrix elements and θ and ϕ are the polar and azimuthal angles for the orientation of the rotating axes of the nitroxide bonded to the lipid relative to the director of the bilayer, i.e. the preferential orientation of lipid molecules; the angular brackets imply ensemble averaging. S_0_ and its uncertainty were then calculated in well-known fashion from its definition and the dimensionless ordering potentials C_20_ and C_22_ and their uncertainties found in the fitting. The typical uncertainties we find for S_0_ range from 1-5 × 10^−3^, while the uncertainties from repeated experiments are 5-8 × 10^−3^ or less than ±0.01. S_0_ indicates how strongly the chain segment to which the nitroxide is attached is aligned along the normal to the lipid bilayer, which is strongly correlated with hydration/dehydration of the lipid bilayers [31,47].

